# Rapid evolution of seed dormancy during sunflower de-domestication

**DOI:** 10.1101/2021.05.21.445131

**Authors:** Fernando Hernández, Roman B. Vercellino, Claudio Pandolfo, Jennifer R. Mandel, Alejandro Presotto

## Abstract

Hybridization between crops and their wild relatives may promote the evolution of de-domesticated (feral) weeds. Wild sunflower (*Helianthus annuus* L.) is typically found in ruderal environments, but crop-wild hybridization may facilitate the evolution of weedy populations. Using one crop-specific mitochondrial marker (CMS-PET1) and 14 nuclear SSR markers, we studied the origin and genetic diversity of a recently discovered weedy population of sunflower (named BRW). Then, using a resurrection approach, we tested for rapid evolution of weedy traits (seed dormancy, herbicide resistance, and competitive ability) by sampling weedy and wild populations 10 years apart (2007 and 2017). All the weedy plants present the CMS-PET1 cytotype, confirming their feral origin. At the nuclear markers, BRW showed higher genetic diversity than the cultivated lines and low differentiation with one wild population, suggesting that wild hybridization increased their genetic diversity. We found support for rapid evolution towards higher seed dormancy, but not for higher competitive ability or herbicide resistance. Our results highlight the importance of seed dormancy during the earliest stages of adaptation and show that crop-wild hybrids can evolve quickly in agricultural environments.

## Introduction

Since the origins of agriculture, farmers have grown domesticated plants in artificially benign environments, characterized by frequent disturbance, high nutrient and water availability, artificially controlled pests, and monoculture or short-term rotations (Clement 2014). These modified environments are beneficial for maximizing crop yields, but they also represent novel niches for opportunistic plant species, namely agricultural weeds (Baker 1974; De Wet and Harlan 1975). Here, we define weeds (or weedy populations) as those populations that interfere with agriculture, regardless of their wild, cultivated, or mixed origin, and wild (or ruderal) populations as those that grow and reproduce away from the agriculture environments (outside agricultural plots), whether native or not (Ellstrand *et al*. 2010). Weeds compete with crops for resources, such as light, nutrients, and water, representing a major cause of crop losses worldwide, estimated at 34 % reduction in crop productivity (Oerke 2006).

Agricultural environments exert high selection pressures on weeds, and weeds often respond by evolving a suite of traits that increase fitness in these environments, such as seed dormancy, herbicide resistance, early flowering, rapid growth, and high reproductive output (Baker 1974; Fried *et al*. 2012; Huang *et al*. 2017). However, is not clear what traits are instrumental for the successful establishment of weeds and what traits are selected later, once weeds are established. Identifying traits associated with the earliest stages of adaptation to agricultural environments is critical to better understand the emergence of novel weedy populations and to design better weed management strategies. Seed dormancy plays a crucial ecological role, contributing to avoidance of out of season germination (Presotto *et al*. 2020; Hernández *et al*. 2021), and persistence in soil seed banks (Darmency *et al*. 2017; Presotto *et al*. 2020). On the other hand, the high killing rates associated with herbicide applications exert strong selection pressure for herbicide resistance (Tranel and Wright 2002; Darmency *et al*. 2017; Kreiner *et al*. 2018).

Herbicides inhibiting the enzyme acetohydroxy acid synthase (AHAS) are among the most widely used herbicides worldwide (Tan *et al*. 2005). The simple genetic architecture of AHAS resistance, and the discovery of alleles confering resistance in several cultivated species, allowed the development of the non-transgenic Clearfield® technology, which confers AHAS resistance to several crop species, including wheat, rice, sunflower, and oilseed rape (Tan *et al*. 2005). However, the drawback of this group of herbicides is their propensity to generate resistant biotypes (Tranel and Wright 2002). AHAS resistant weeds have been reported in at least 166 species worldwide (Tranel *et al*., 2021), including weedy sunflower (Al-Khatib *et al*. 1998; White *et al*. 2002).

Weeds can originate in three ways: 1) from crop varieties (endoferality); 2) from wild populations; and 3) from crop-wild hybrids (exoferality) (Baker 1974; Ellstrand *et al*. 2010; Huang *et al*. 2017). Ferality or de-domestication is an evolutionary process by which domesticated species escape from farmers’ fields and acquire the ability to form self-perpetuating populations (Gressel 2005). Then, when these biotypes invade agricultural fields, they are designated as feral weeds (Gressel 2005; Ellstrand *et al*. 2010). Regardless of weeds origin, weedy traits may evolve by selection on the standing genetic variation (Huang *et al*. 2017), new mutations (Al-Khatib *et al*. 1998; Vercellino *et al*. 2018), or by adaptive introgression (Pandolfo *et al*. 2018; Le Corre *et al*. 2020), defined as the permanent incorporation of foreign variants through backcrossing, and mantained by natural selection (Suarez-Gonzalez *et al*. 2018).

A powerful technique for studying rapid evolution is the “resurrection approach” (Franks *et al*. 2018). This approach consists in reviving ancestors from stored propagules (often dormant seeds) and comparing them with descendants under common conditions. In this way it is possible to evaluate the phenotypic and genetic changes in response to the most important selective forces (e.g., drought stress, or global warming) that may have occurred during the time between collections (Kuester *et al*. 2016; Franks *et al*. 2018; Weis 2018). This approach has been used to study rapid phenotypic and/or genetic changes in plants in response to drought (Franks *et al*. 2016), and herbicide use (Kuester *et al*. 2016), demonstrating that the evolution of adaptive traits may occur in a few generations (< 10) if the selective forces are strong enough.

Wild sunflower (*Helianthus annuus* L.) is native to North America and introduced in many areas, including southern Europe (Muller *et al*. 2011) and Argentina (Poverene *et al*. 2008). It is a typical ruderal species, growing in roadsides, field margins and ditches, although weedy populations have evolved both in the native (Kane and Rieseberg 2008; Mayrose *et al*. 2011) and introduced areas (Muller *et al*. 2011; Casquero *et al*. 2013). In the native area, the repeated use of herbicides in agricultural fields has resulted in the evolution of sunflower populations resistant to AHAS herbicides (Al-Khatib *et al*. 1998; White *et al*. 2002), and more recently to glyphosate (Singh *et al*. 2020).

In Argentina, wild sunflower was introduced at least 70 years ago, probably with breeding purposes, and has subsequently spread over the central region (Poverene *et al*. 2008). Argentinean populations are mainly found in ruderal environments and show genetic and phenotypic similarity to native wild populations from North America (Hernández, Presotto, *et al*. 2019). In the country, several hybrid zones were formed, where wild and cultivated sunflower often grow in sympatry and natural hybrids can be seen (Ureta *et al*. 2008; Mondon *et al*. 2018). However, a previous study found evidence of crop to wild introgression in Argentinean populations of prairie sunflower (*H. petiolaris*) but not in *H. annuus* (Mondon *et al*. 2018), suggesting that ecological barriers (e.g., low fitness of hybrids) are preventing the establishment of crop-wild hybrids in hybrid zones. In 2003, sunflower cultivars with the Clearfield® technology, which confers AHAS resistance, were introduced and then rapidly adopted by Argentinean farmers, heightening the risk of herbicide resistance evolution through crop to wild gene flow (Presotto *et al*. 2012).

In 2007, a population of *H. annuus* was found in Barrow (hereafter BRW), near Tres Arroyos, Buenos Aires, a region with no previous records of wild *Helianthus* species (Casquero *et al*. 2013). Such population is the first reported as an agricultural weed in Argentina, infesting sunflower and maize crops (Casquero *et al*. 2013). This weed showed intermediate traits between crop and wild, suggesting it is a natural crop-wild hybrid (Casquero *et al*. 2013; Presotto *et al*. 2017). A previous study showed that when glyphosate-resistant soybean was planted, no weedy sunflower plants were observed but when sunflower or maize were planted, up to 48,000 weedy sunflowers per ha^-1^ were found (Casquero *et al*. 2013), suggesting that chemical control in these crops is not effective. Given that chemical control of weeds in sunflower and maize mostly relied on AHAS-inhibiting herbicides, we hypothesize that AHAS resistance has contributed to the adaptation of weedy sunflower in the agricultural environment.

Here, we studied the origin, genetic diversity and phenotypic evolution of BRW. First, we compared BRW with wild and cultivated sunflower using a crop-specific marker (CMS-PET1) and 14 nuclear SSR markers. Then, using a resurrection approach, we tested for rapid evolution of weedy traits (seed dormancy, herbicide resistance, and competitive ability) in response to agricultural conditions by sampling BRW 10 years apart (2007 and 2017). The specific aims were to 1) investigate the origin and genetic diversity of BRW, and 2) test whether BRW shows evidence of rapid evolution of weedy traits.

## Materials and methods

### Plant material

We focused our study on BRW, a weedy population collected near Tres Arroyos, Buenos Aires (Fig. 1; Table S1). The agricultural field where BRW was collected is managed by farmers following a summer crop rotation system, alternating sunflower, glyphosate-resistant soybean, and maize (Casquero *et al*. 2013). For BRW, an estimated population size of up to 4.8 × 10^5^ were registered in four successive years (Casquero *et al*. 2013).

**Figure 1.**
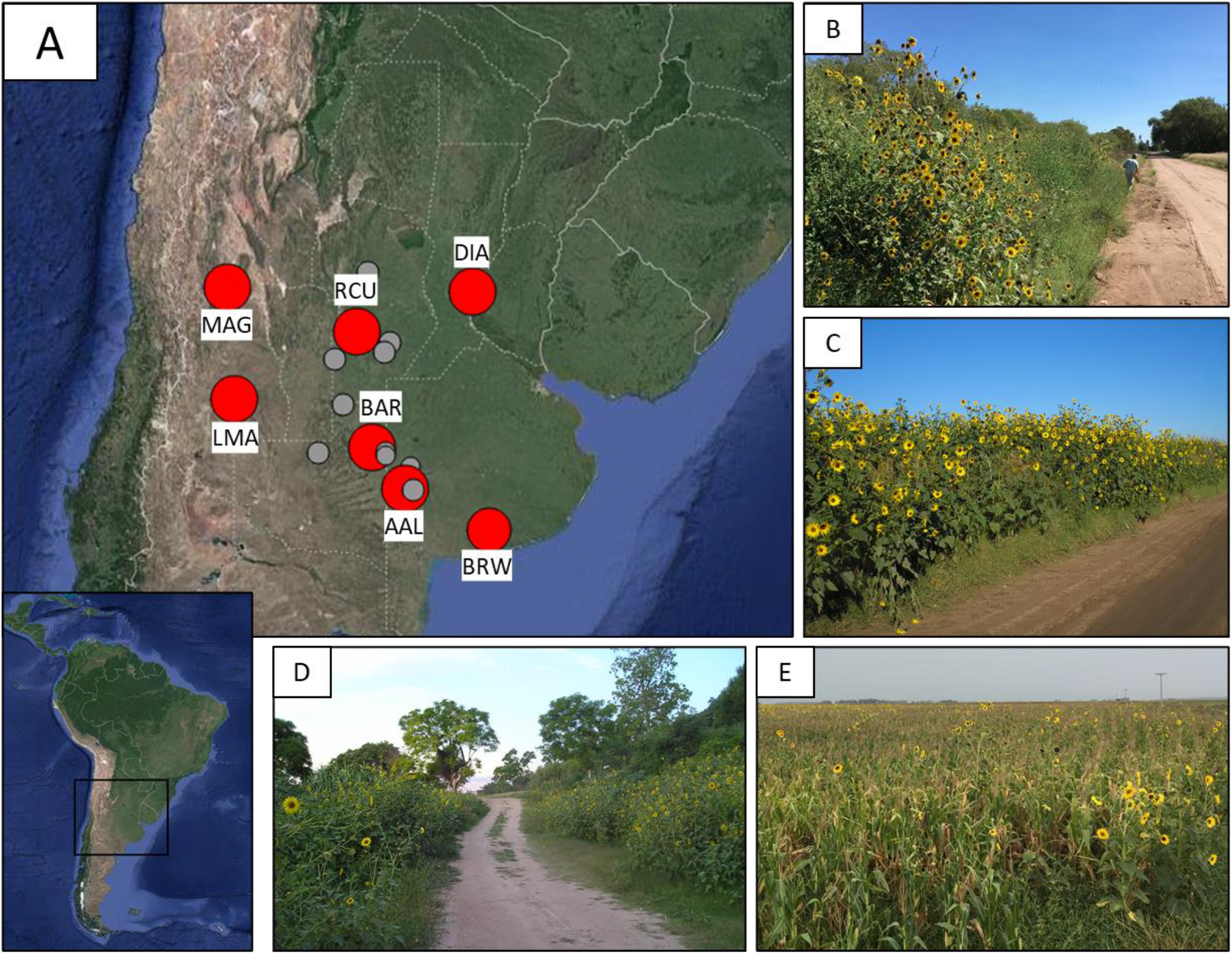
Geographical location of *Helianthus annuus* populations in Argentina (A), red dots for populations used in this study, grey dots for populations not used in this study included to show the overall distribution of wild *H. annuus* in Argentina. B-D: ruderal populations from Colonia Baron (BAR; B), Rio Cuarto (RCU; C), and Diamante (DIA; D). E: weedy population growing in a maize plot near Tres Arroyos (BRW). Pictures from the collection trip in March 2017. See online version for full color.

### Populations sampled for genotyping and population genetic analysis

In cultivated sunflower, hybrid seed production is based on the CMS-PET1 mitochondrial cytotype, discovered by Leclercq (1969) and a few Rf nuclear genes. The CMS-PET1 cytotype is present in the female parents to ensure cross-pollination while Rf genes are carried by male parents in homozygosis to restore pollen fertility in the hybrid progeny. As the cytoplasm is maternally inherited, all cultivars carry the CMS-PET1 cytotype, whereas it is absent in wild sunflowers (Rieseberg *et al*. 1994). Thus, markers differentiating CMS-PET1 from non-CMS cytotypes can be used to infer the cultivated or wild maternal origin of weeds (Rieseberg *et al*. 1994; Muller *et al*. 2011; Garayalde *et al*. 2015).

We used the CMS-PET1 marker to discern the cultivated or wild origin of BRW. We genotyped 32 samples from Argentinean populations (8 individuals each for BAR, BRW, DIA, and RCU), two commercial cultivars grown in Argentina (Cacique CL and HS03) as positive controls, and 10 samples from North American populations: three individuals from North Dakota (ND; PI 586888) and Texas (TX; PI 613728), and four individuals from California (CA; PI 413131). To detect the occurrence of the CMS-PET1 cytotype, we used a duplex PCR strategy, *orfC* primers were used to indicate the presence/absence of the CMS-PET1 cytotype, and *coxIII* was included as a positive control (Rieseberg *et al*. 1994; Garayalde *et al*. 2015). Primer sequences, PCR conditions, and the expected size of PCR products for *orfC* and *coxIII* can be found in Garayalde *et al*. (2015).

To study the genetic diversity and population structure, we collected DNA from 79 individuals, 60 belonging to six ruderal populations collected in central Argentina (Hernández, Poverene, *et al*. 2019), plus 10 individuals from BRW, and nine cultivated inbred lines from different origins (Table 1). Despite their origin, all cultivated lines were widely used in Argentina for germplasm development (Filippi *et al*. 2015). Individuals were genotyped using 14 nuclear microsatellites, a detailed description of the DNA extraction and SSR genotyping protocols can be found in Hernández, Presotto, *et al*. (2019). Three individuals (one from BRW and two from RCU) with missing data at more than three loci were removed from further analyses.

**Table 1.**
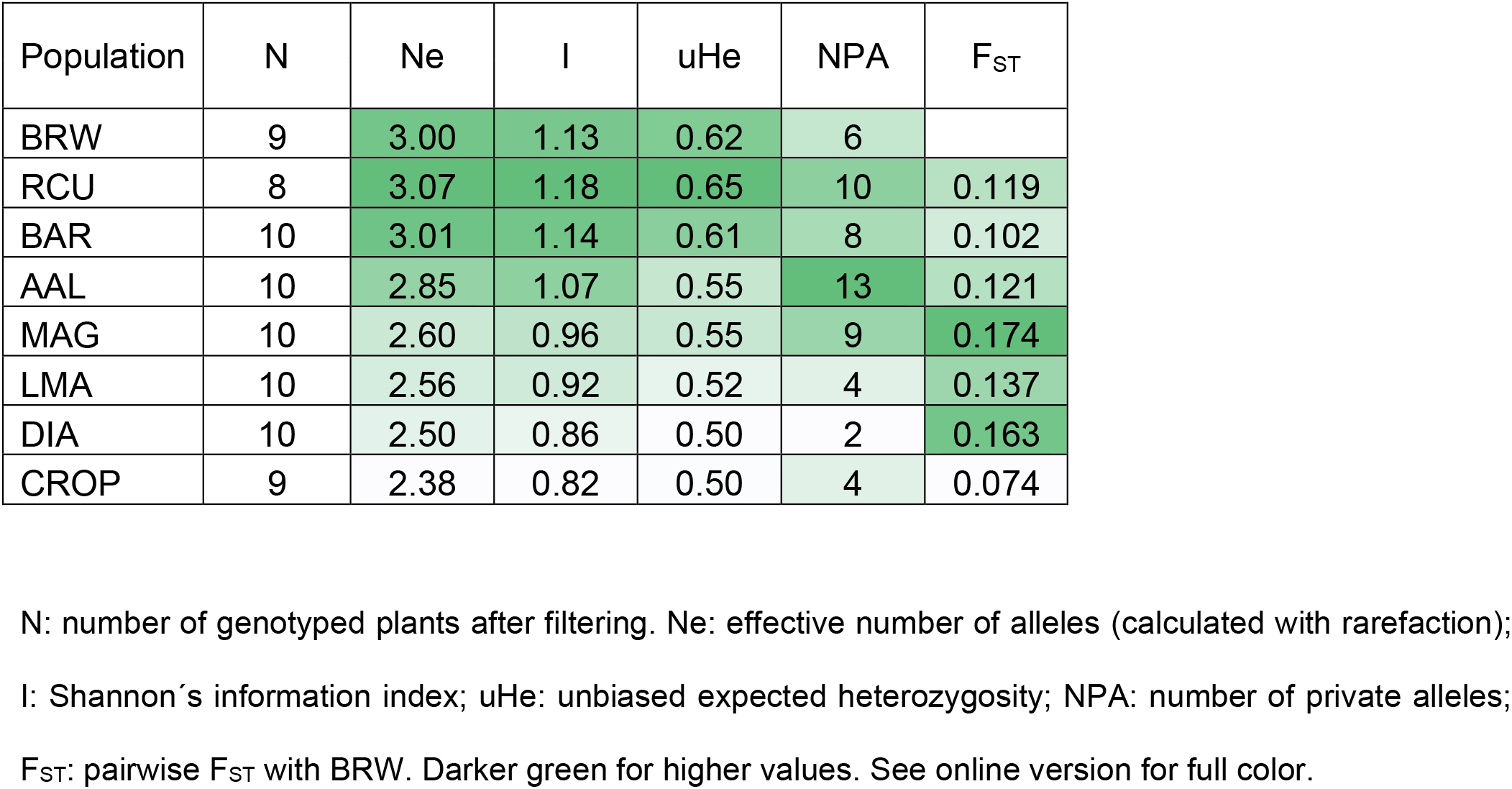
Measurements of genetic diversity per population.

| Population | N | Ne | I | uHe | NPA | F<sub>ST</sub> |
| --- | --- | --- | --- | --- | --- | --- |
| BRW | 9 | 3.00 | 1.13 | 0.62 | 6 |  |
| RCU | 8 | 3.07 | 1.18 | 0.65 | 10 | 0.119 |
| BAR | 10 | 3.01 | 1.14 | 0.61 | 8 | 0.102 |
| AAL | 10 | 2.85 | 1.07 | 0.55 | 13 | 0.121 |
| MAG | 10 | 2.60 | 0.96 | 0.55 | 9 | 0.174 |
| LMA | 10 | 2.56 | 0.92 | 0.52 | 4 | 0.137 |
| DIA | 10 | 2.50 | 0.86 | 0.50 | 2 | 0.163 |
| CROP | 9 | 2.38 | 0.82 | 0.50 | 4 | 0.074 |

N: number of genotyped plants after filtering. Ne: effective number of alleles (calculated with rarefaction);
I: Shannon's information index; uHe: unbiased expected heterozygosity; NPA: number of private alleles;
F<sub>ST</sub>: pairwise F<sub>ST</sub> with BRW. Darker green for higher values. See online version for full color.

The effective number of alleles per population was calculated with rarefaction using the *hierfstat* R package (Goudet 2005), the Shannon’s information index and unbiased gene diversity were calculated at the population level using GenAlEx v. 6.5 (Peakall and Smouse 2012). Population structure was assessed using analysis of molecular variance (AMOVA), as implemented in GenAlEx. Bayesian clustering, using the STRUCTURE v.2.3.4 software (Pritchard *et al*. 2000) was used to assign multi-locus genotypes into clusters, as in Hernández, Poverene, *et al*. (2019). The population structure was also studied using discriminant analysis of principal components (DAPC), with the *adegenet* R package (Jombart *et al*. 2010).

### Resurrection experiments

For the resurrection experiments, the weedy BRW and three ruderal populations (BAR, DIA, and RCU) were sampled 10 years apart (2007 and 2017). Ruderal populations were included to test for evolution due to conditions other than the agricultural ones (e.g., due to global warming or genetic drift). Population size of both ancestor and descendant populations were large (> 10,000 individuals). For sampling, we followed the recommendations of Franks *et al*. (2018). Seeds from > 100 plants were collected at the same site as inferred using GPS coordinates (Fig. 1; Table S1). To reduce the so called “invisible fraction bias” (Weis 2018), consisting in mean phenotypic variation due to non-random mortality during storage, the achenes (hereafter seeds) for the ancestral populations (2007) were regenerated once (in 2011) in a common garden through controlled pollination of 25 - 50 individuals.

### Germination experiments

For the gemination experiments, we used the four populations sampled 10 years apart (2007 and 2017): BAR, BRW, DIA, and RCU (Fig. 1). Two germination experiments were conducted in two different years. For experiment 1 (2016/17), seeds of the ancestral populations (2007), regenerated once in 2011, were produced in a common garden at the Agronomy Department, Universidad Nacional del Sur, Bahia Blanca, Argentina (38°41’38’’S, 62°14’53’’W), whereas for descendant populations (2017), seeds were collected from more than 100 plants per population from their local environments in a three-day collection trip. For experiment 2 (2017/18) all the seeds were produced in a common garden at the Agronomy Department, Universidad Nacional del Sur, to control for maternal environmental effects. In common gardens, seeds for each population were produced under controlled pollination of the heads of 15–20 plants covered with paper bags at the pre-flowering stage, following Hernández *et al*. (2017). At the flowering stage, heads were hand-pollinated with pollen from sibling plants, and seeds from these 15-20 plants (one head per plant) were pooled and used for germination experiments.

The germination experiments were carried out following Hernández, Poverene, *et al*. (2019). Briefly, germination was evaluated at two different times: two months (T1) and eight months (T2) after harvest, and at three constant temperatures (10 °C, 20 °C, and 30 °C). After harvest, seeds were stored in tri-laminar aluminum bags to protect the seeds from humidity, at room temperature (25 °C) for 2 months (T1). Then, seeds were placed in a growth chamber at 5 °C for six months (T2). Four replicates of 25 seeds per population and treatment were placed on filter paper in petri dishes and moistened with distilled water, with a 12 h photoperiod and they were counted periodically at 2–3 days intervals for 16 days. Replicates were arranged in four complete blocks (a rack of the growth chamber). For each experiment, the germination proportion (GRP) was calculated as the ratio between germinated seeds at the end of the experiment (day 16) and viable seeds. Seed viability of was obtained using a tetrazolium test on non-germinated seeds for all the replicates at the end of each experiment (ISTA, 2004). Viable seeds that did not germinate were considered dormant seeds.

Data were analyzed with generalized linear models using PROC GLM in SAS (SAS University edition; SAS Institute Inc., Cary, NC). Each experiment and time were analyzed separately resulting in four analyses. The single effects were the collection Year (2007 and 2017), Population (BAR, BRW, DIA, and RCU), and Temperature (10, 20, and 30°C), besides the two-way and three-way interactions. Due to the high homogeneity in the blocks (tested in preliminary analyses) we did not include the block effect in the final analyses. All effects were considered as fixed. When the Year*Population interaction was significant, the least square means of each population were compared between years, and when the Year*Population*Temperature interaction was significant, the least square means of each population and temperature were compared between years. Significance values were adjusted for multiple comparisons with the Tukey-Kramer test. Due to the lack of seeds, some populations were not evaluated at all three temperatures in all the experiment and time combinations.

### Competition experiment

To evaluate whether higher competitive response evolved in BRW, we performed a target-neighborhood design. Three sunflower populations at three levels of maize competition were evaluated in a factorial experimental design with five replicates. Competition levels consisted in 0, 1, and 3 maize plants per pot (Treatments 1, 2, and 3, respectively), and populations were BAR (the putative wild parent of BRW, collected in 2017), and BRW sampled in 2007 (hereafter BRW_07_) and 2017 (hereafter BRW_17_). For Treatments 2 and 3, the maize plants were established by sowing one and three seeds, respectively, in 15 cm diameter plastic pots (2.5 L) while no maize seeds were sown for the Treatment 1. At the two-leaf stage of maize, sunflower plants were established by sowing one pre-germinated sunflower seed per pot. Sunflower was planted after maize establishment to ensure interspecific competition. The experiment was carried out in a greenhouse (25 °C ± 3 °C), with light provided by six 36 W fluorescent tubes located 50 cm above plants, and the pots were watered manually to keep them at field capacity throughout the experiment. Three variables (plant height, leaf width and leaf length) were measured four times on sunflower plants on 13, 17, 22, and 30 days after sowing, and the leaf size was estimated as leaf width*leaf length for each time. At the end of the experiment (30 days after sowing), the aboveground biomass of sunflower plants was collected by cutting the entire plants at ground level, drying the tissue at 60 °C for 7 days, and weighing it.

We performed a multivariate analysis of variance (MANOVA) on the 17 variables (four variables measured at four times plus aboveground biomass at the end of the experiment) using PROC GLM in SAS. Fixed effects were competition Treatment (three levels), Population (BAR, BRW_07_, and BRW_17_), and the Treatment*Population interaction. Significance was evaluated using the Wilks’ Lambda test criterion.

### Herbicide resistance experiment

To evaluate herbicide resistance in BRW, the response of BRW_07_ and BRW_17_ to imazapyr (an AHAS-inhibiting herbicide) was evaluated, following Vercellino *et al*. (2018). The progeny of one herbicide resistant cultivar (Cacique CL, hereafter IMI) and a known susceptible population (BAR) were included as resistant and susceptible controls, respectively. For the screening, plants were established by sowing pre-germinated seeds in 200-cell plastic trays (50 cc per cell) in a complete randomized design with three replicates of 17 (BAR and IMI) or 45 (BRW_07_ and BRW_07_) plants, totaling 372 plants. At the two-leaf stage, Imazapyr was applied at 1.2 X the recommended rate (X = 210 mL ha^-1^) and the survival rate was recorded 18 and 35 days after herbicide application. Based on survival rate, populations were qualitatively classified as resistant or susceptible to the herbicide. Plants were grown in a greenhouse at 20-25 °C, from sowing to the end of the experiment.

## Results

### Origin and genetic diversity of the weedy population

Cytoplasmic analysis shows that CMS-PET1 cytotype was present in all the individuals of BRW (n = 8), but it was absent in all wild individuals from either Argentina (n = 29) or the United States (n = 10), indicating a feral origin of BRW.

To find out whether the origin of BRW is endoferal (from a crop escape) or exoferal (resulting from crop-wild hybridization), we analyzed the genetic diversity and population structure of BRW along with wild populations and cultivated lines (CROP), using 14 nuclear SSR markers. If BRW is endoferal, its genetic diversity should be similar or lower than genetic diversity observed in CROP and BRW samples should group with CROP. If BRW is exoferal, its genetic diversity should be higher than CROP and BRW samples should be intermediate between cultivated and wild samples in population genetic approaches.

In terms of genetic diversity, BRW exhibited at least 20% more genetic diversity than CROP, and similar values to RCU and BAR for any measure (Table 1), suggesting that wild hybridization increased the genetic diversity of the feral BRW.

Population structure was evaluated using three independent approaches (pairwise F_ST_, STRUCTURE, and DAPC). The pairwise F_ST_ between BRW and the other populations ranged from 0.07 (BRW-CROP) to 0.17 (BRW-MAG) (Table 1). The lowest F_ST_ values were observed with CROP and BAR, whereas the highest were with DIA and MAG (Table 1). STRUCTURE analysis suggests three, followed by five genetic groups (Fig. 2A, S1A). At K=3, individuals of BRW grouped with CROP and DIA individuals (Fig. 2A). At K=5, individuals of BRW showed evidence of an admixture between BAR and CROP (Fig. 2A). DAPC assigned individuals into four groups (Fig. 2B, S1B), and group 2 included all the individuals from BRW and most (eight out of nine) cultivated lines and BAR individuals (seven out of ten) (Fig. 2C).

**Figure 2.**
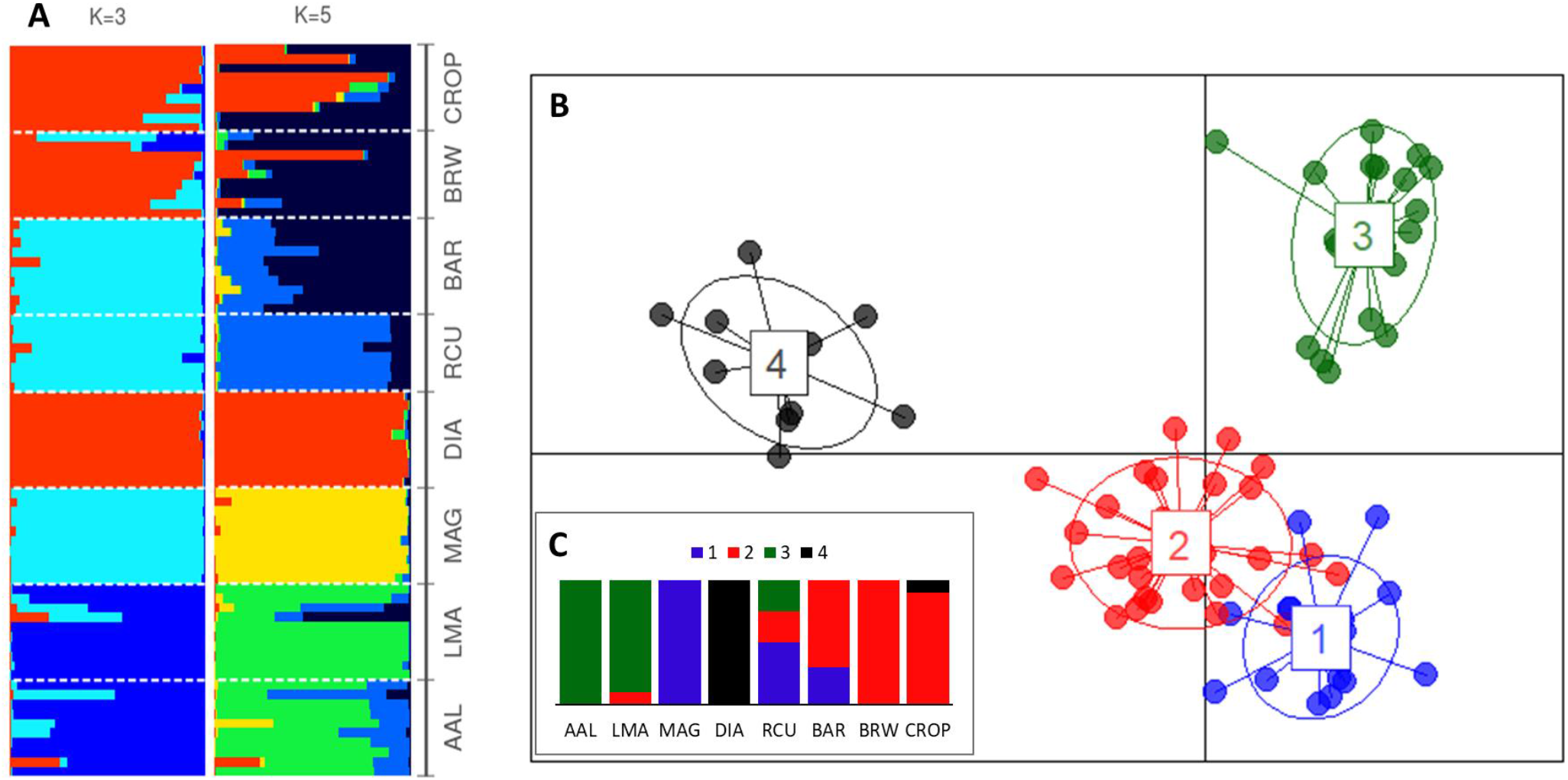
Population structure of wild, cultivated, and weedy sunflower. A: STRUCTURE output; B: DAPC output; C: proportion of individuals assigned to each DAPC group. Wild populations: AAL, BAR, DIA, LMA, MAG, and RCU; weedy: BRW; cultivated: CROP. Details of populations in Table S1. See online version for full color.

### Germination experiments

To test for phenotypic evolution between the years of collection, we explored the Year*Population and Year*Population*Temperature interactions.

We observed significant Year*Population*Temperature interactions in all four experiment and time combinations (P = 0.06 for Experiment 1 and Time1, P < 0.01 for the rest), indicating that some populations differ between collection years, and these responses varied with temperature. In Experiment 1, BRW_17_ showed a significantly lower germination proportion (GRP) than BRW_07_ in all six experimental combinations (Fig. 3A, 3B), whereas DIA and RCU collected in 2017 had a lower GRP than those collected in 2007, but only at 20 °C (Fig. 3A, 3B). In addition, at Time 1 and 10°C, DIA_17_ showed a higher GRP than DIA_07_ (Fig. 3A). Similar results were observed in experiment 2, when all the seeds were produced in a common garden. BRW_17_ showed a significantly lower GRP than BRW_07_ in all four combinations with some germination, whereas DIA_17_ and RCU_17_ showed a lower GRP than DIA_07_ and RCU_07_ at 20°C (Fig. 3C, 3D). We also observed significantly lower GRP in DIA_17_ than in DIA_07_ at 10°C (Fig. 3C, 3D).

**Figure 3.**
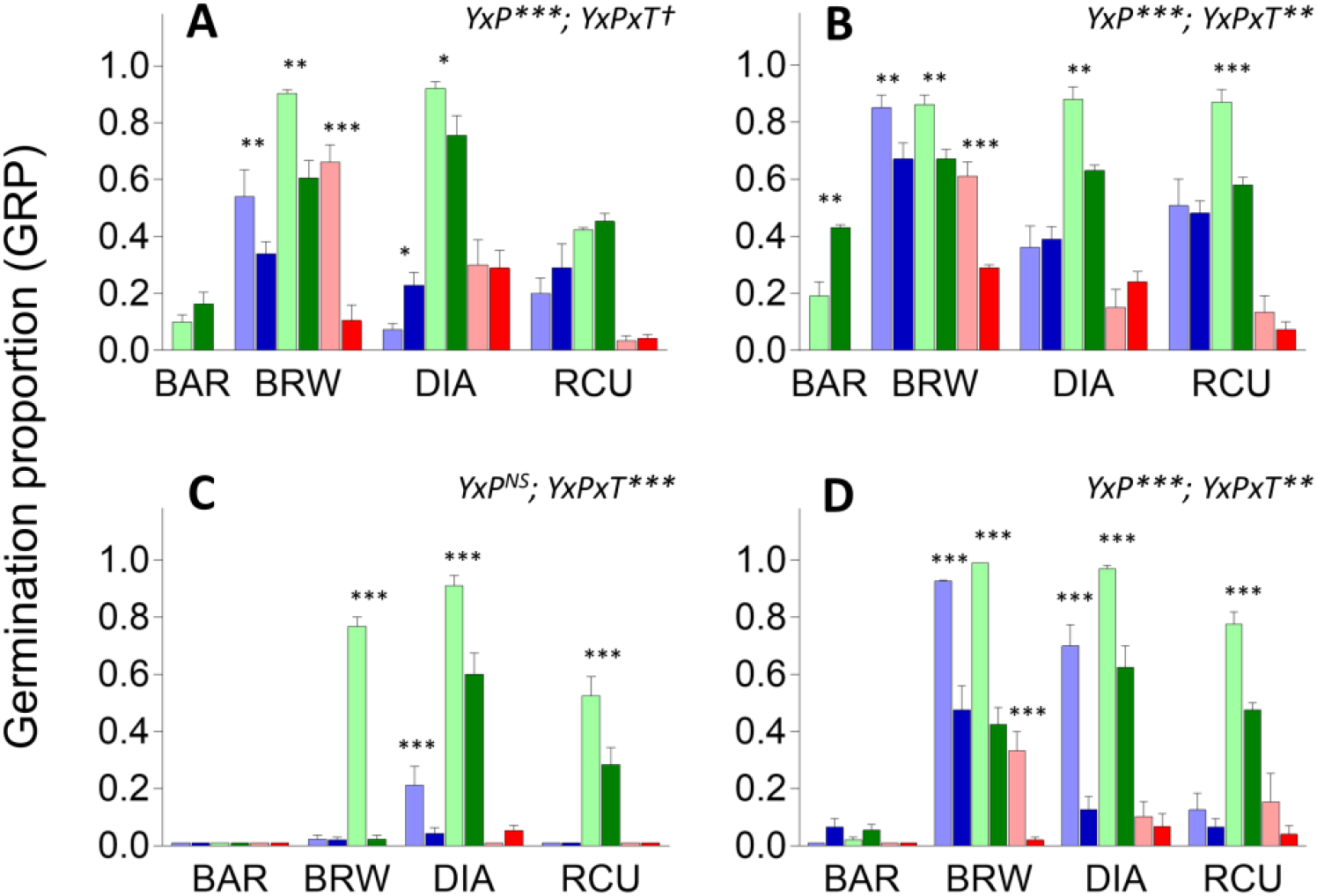
Germination proportion (GRP) measured in two times over two experiments at 10, 20, and 30°C, indicated by blue, green, and red color bars, respectively. A: experiment 1, time 1; B: experiment 1, time 2; C: experiment 2, time 1; D: experiment 2, time 2. Within each temperature lighter bars are used for populations collected in 2007 and darker bars for populations collected in 2017. Significance of Year*Population (Y*P) and Year*Population*Temperature (Y*P*T) interactions are indicated in the upper right of each plot. When Y*P*T is significant, least squares means of populations collected in 2007 and 2017 are compared within each temperature. *** P < 0.0001; ** P < 0.01; * P < 0.05; † P < 0.1; NS P > 0.1. See online version for full color.

### Interspecific competitive ability

In the MANOVA, we observed significant Population and Treatment effects and significant Population*Treatment interaction. To unravel this interaction, we compared populations within treatments by using *a priori* orthogonal contrasts (Table 2). Without maize competition (Treatment 1), all three populations showed significant differences (Table 2) and for most traits BRW_17_ showed larger values than BRW_07_, followed by BAR (Fig. 4). With maize competition (Treatments 2 and 3), BAR showed significant differences with both BRW_07_ and BRW_17_, but no significant differences were found between BRW_07_ and BRW_17_ (Table 2) and for most traits, BRW_17_ and BRW_07_ showed larger values than BAR (Fig. 4).

**Table 2.** Multivariate analysis of variance of 17 phenotypic traits measured in the competition experiment. Significant contrasts in bold.

| Effect | F | P |
| --- | --- | --- |
| Population | 6.61 | < 0.0001 |
| Treatment | 5.00 | < 0.0001 |
| Population*Treatment | 1.68 | 0.0197 |
| Contrasts |  |  |
| T1 BAR vs BRW<sub>07</sub> | 4.9 | **0.0017** |
| BAR vs BRW<sub>17</sub> | 10.36 | **< 0.0001** |
| BRW<sub>07</sub> vs BRW<sub>17</sub> | 4.97 | **0.0016** |
| T2 BAR vs BRW<sub>07</sub> | 3.5 | **0.0094** |
| BAR vs BRW<sub>17</sub> | 4.29 | **0.0034** |
| BRW<sub>07</sub> vs BRW<sub>17</sub> | 0.74 | 0.7309 |
| T3 BAR vs BRW<sub>07</sub> | 2.49 | **0.0415** |
| BAR vs BRW<sub>17</sub> | 3.06 | **0.0175** |
| BRW<sub>07</sub> vs BRW<sub>17</sub> | 1.57 | 0.1917 |

**Figure 4.**
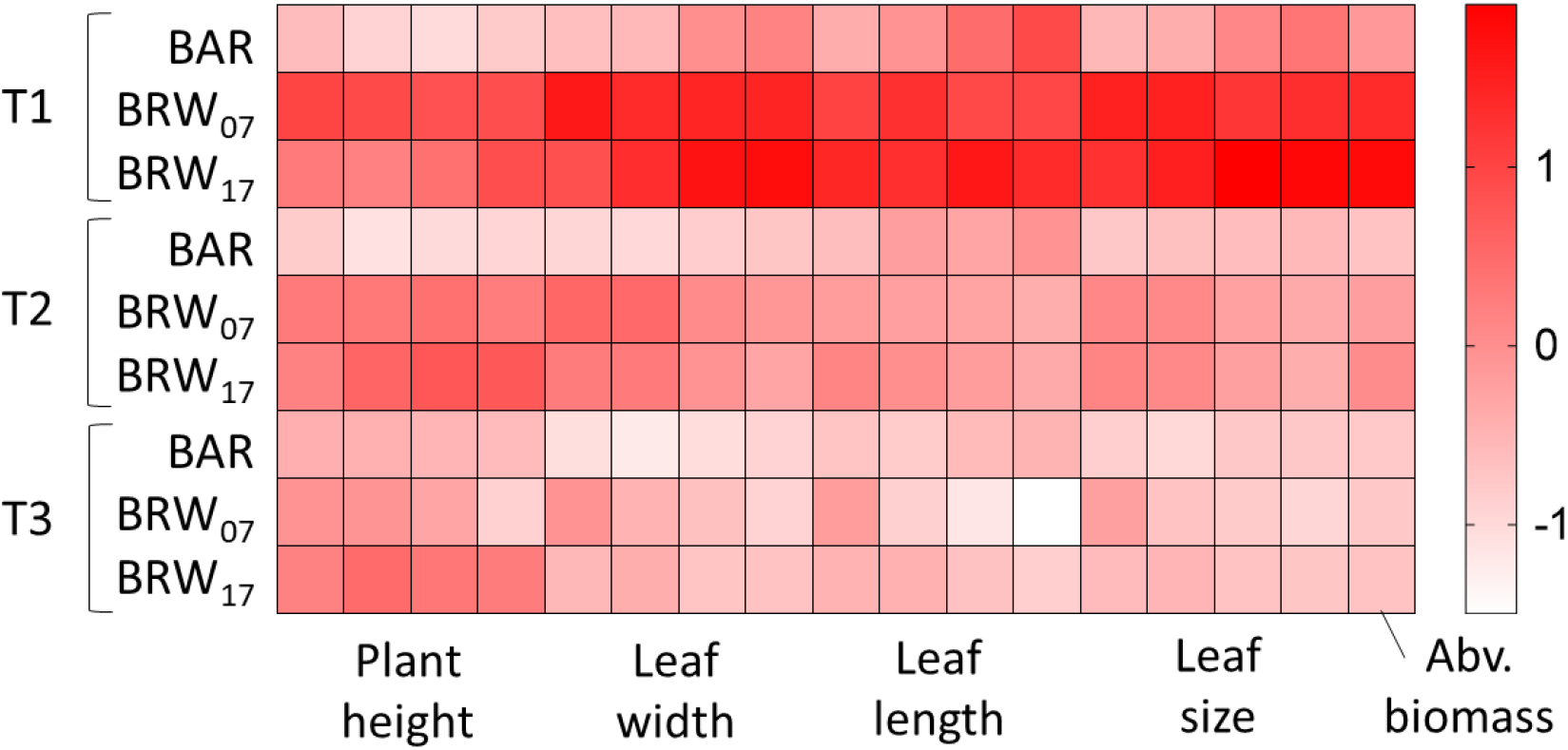
Heatmap of 17 standardized phenotypic variables (mean = 0, SD = 1) of three populations (BAR, BRW_07_, and BRW_17_) at three competition treatments (T1 - T3). Shading intensity increase with larger phenotypic values. See online version for full color.

### Herbicide resistance

At a 1.2 X the recommended rate of imazapyr and 35 days after herbicide application, plant survival was null for BAR (0/46) and BRW_07_ (0/126) and very low for BRW_17_ (1/129), whereas it was high for the progeny of the herbicide resistant cultivar (IMI) (41/48).

## Discussion

### On the origin of BRW

Understanding the origin and evolution of agricultural weeds is critical for designing better weed management strategies, especially for preventing the emergence of novel biotypes and the escape of genes/alleles from cultivated species. Our results, based on the CMS-PET1 confirmed the feral origin of BRW proposed by Casquero *et al*. (2013). All individuals of BRW harbor the CMS-PET1 cytotype, used in commercial hybrid seed production (Rieseberg *et al*. 1994; Muller *et al*. 2011). The CMS cytotype has previously been used to confirm the feral origin of weedy sunflower populations in Europe (Muller *et al*. 2011). In addition, all three population genetic approaches we employed (F_ST_, STRUCTURE, and DAPC) grouped BRW with CROP, supporting a feral origin. We also found that BRW shows greater genetic diversity than CROP, with a diversity as high as the wild populations, and group with one wild population, supporting a crop-wild origin.

The weedy BRW was collected from a region with no previous records of wild sunflower (Poverene *et al*. 2008), thus, where did hybridization take place? Hybrid zones, where crop and wild relatives grow in sympatry are potential sources of crop-wild hybrids. Two of our sampled populations often grow in sympatry with the crop: AAL in Adolfo Alsina, and BAR in Colonia Baron, and natural crop-wild hybrids are often observed there (Ureta *et al*. 2008; Mondon *et al*. 2018). In Argentina, the genetic diversity of wild sunflower is geographically structured (Hernández, Presotto, *et al*. 2019), making population genetic approaches useful for identifying the geographic source of BRW. Of the two hybrid zones, BAR is the most probable source. All the approaches we used (pairwise F_ST_, and clustering analyses) showed that BAR was the wild population that was genetically closest to BRW. Surprisingly, we found some genetic similarity between DIA and BRW, especially in the STRUCTURE clustering. Although the origin of DIA is unclear (Hernández, Presotto, *et al*. 2019), this population does not grow in sympatry with cultivated sunflower (Poverene *et al*. 2008). A probable origin of DIA is the intentional introduction by European migrants during early 1900’s (Hernández, Presotto, *et al*. 2019), therefore, it is possible that both BRW and DIA share a feral origin: BRW from modern cultivars (all plants harbor the CMS-PET1 cytotype), and DIA from landraces or open pollinated varieties (no plants harbor the CMS-PET1 cytotype). On the other hand, the farmer who manages the field where BRW was collected indicated that the first plants of multiheaded sunflower were seen between 2003 and 2005 after renting a harvest machine from an area close to Colonia Baron, where BAR is naturalized. As no wild plants were observed in BRW but crop-wild hybrids are often observed in BAR (Ureta *et al*. 2008; Mondon *et al*. 2018), we hypothesize that hybridization occurred in Colonia Baron, and then hybrid seeds were accidentally introduced into Barrow.

Feral weeds have been reported in a dozen cultivated species (reviewed in Ellstrand *et al*. 2010). In sunflower, feral forms have been found in agricultural fields across southern Europe, where the wild species is absent (Muller *et al*. 2011). There are three similar features between BRW and European populations (Muller *et al*. 2011; Casquero *et al*. 2013; Presotto *et al*. 2017): 1) the exoferal origin, in both cases natural crop-wild hybrids were accidentally introduced to farmers′ fields; 2) allopatry with wild sunflower, gene flow with wild sunflower was disrupted when crop-wild hybrids were introduced to a novel region; and 3) the adaptation to agricultural environments, an ecological transition for a typically ruderal species (Kane and Rieseberg 2008; Poverene *et al*. 2008). While weedy populations have also evolved from wild ruderals (Kane and Rieseberg 2008; Mayrose *et al*. 2011), crop-wild hybrids are rarely established in ruderal environments (Poverene *et al*. 2008; Mondon *et al*. 2018). This indicates that crop-wild hybridization is not required but it may facilitate the ecological transition from ruderal to agricultural habitats. Recently, we showed that wild-like traits are under selection in both ruderal and agricultural habitats, but selection of wild-like traits is generally weaker in the latter (Presotto *et al*. 2019), making them more permeable for crop to wild introgression In addition, in hybrid zones, gene flow between crop-wild hybrids and wild populations may prevent the adaptation of crop-wild hybrids to agricultural environments, by breaking up favorable allele combinations. This can explain why the hybrids are (apparently) more successful away from hybrid zones. Further studies using natural and synthetic hybrids are needed to directly test the role of hybridization on the evolution in agricultural environments.

### Rapid evolution of seed dormancy

We found strong support for rapid evolution of seed dormancy in BRW, but not for herbicide resistance or competitive ability. BRW_17_ showed much higher dormancy than BRW_07_. Similarly, seed dormancy that has been observed in feral forms of rice (Huang *et al*. 2017), and radish (Vercellino *et al*. 2019) is probably the result of rapid evolution during de-domestication. Due to its crucial ecological role and its early expression in the life cycle, seed dormancy is expected to be under especially strong selection. Recently, we reconstructed the genetic background of BRW by crossing a cultivar (female) with BAR (male) and we observed an overall low seed dormancy with cascading deleterious effects leading to high autumn emergence, low overwinter survival, and poor establishment in the field (Hernández *et al*. 2021). Thus, if genetic variation is present, selection against low dormant phenotypes can be strong.

Evolution from the standing genetic variation is likely to be much faster than from new mutations, especially when multiple small-effect genes are involved (Kreiner *et al*. 2018). In sunflower, no major genes of large effect have been identified for seed dormancy but many small-effect QTLs (Gandhi *et al*. 2005; Brunick 2007), making the evolution of seed dormancy through new mutations in multiple regions very unlikely. Hybridization between wild and cultivated sunflowers leads to high phenotypic and genetic variation on which selection can act (Muller *et al*. 2011; Presotto *et al*. 2019). After hybridization, wild and cultivated alleles starting at high frequencies and novel combinations may produce transgresive phenotypes. In a recent seed bank experiment, we observed that most BRW_07_ seeds germinated in the first year (up to 80%), which is consistent with the low dormant phenotype observed here, but also a small fraction of seeds remained dormant and viable in the soil for at least four years (Presotto *et al*. 2020), indicating that BRW_07_ (our ancestral population) had substantial genetic variation for seed dormancy on which selection could act. In summary, rapid evolution of seed dormancy in BRW was probably facilitated by strong directional selection on the standing genetic variation.

On the other hand, we also found support for rapid evolution of dormancy in ruderals, at least in two populations (DIA and RCU) and one treatment (at 20 °C). To our knowledge, these populations did not experience any major habitat changes since their introduction at least 70 years ago (Poverene *et al*. 2008). As these populations are distributed in patches at least 100 km apart from each other, and harbor thousands of individuals (Poverene *et al*. 2008), it is unlikely that genetic drift and gene flow have played a role in the evolution of seed dormancy. However, from our data, it is not possible to discern whether the evolution of dormancy over the last decade is part of a continous process of local adaptation post-introduction or whether it occured in response to recent habitat changes (e.g., global warming) after populations became locally adapted.

The resurrection approach is one of the most powerful methods for directly testing for rapid evolution (Franks *et al*. 2018; Weis 2018). In our study, we found strong support for rapid evolution towards higher seed dormancy in BRW. Although this approach does not distinguish selection from genetic drift or gene flow (Franks *et al*. 2018), we gathered evidence in favor of selection. First, the phenotype changed in the predicted sense, towards higher seed dormancy (Darmency *et al*. 2017; Presotto *et al*. 2020; Hernández *et al*. 2021). Second, the census size of both ancestor and descendant populations (> 10,000 individuals) and the effective population size of the descendant population (inferred through genetic markers) are very large, minimizing the influence of genetic drift on phenotypic evolution (Kreiner *et al*. 2018). Third, although gene flow between weedy and cultivated sunflower probably occurred in the time frame between the collections, the cultivated sunflower showed mostly no, or low, dormancy (Hernández *et al*. 2017, 2021) and the weedy traits changed in the opposite direction, i.e. dormancy evolved despite, rather than aided by, gene flow. Further studies combining the resurrection approach and population genomics can be used to explore genome-wide changes in allele frequency, scan the genome for signatures of recent selection, and to explore the genetic basis of seed dormancy in weedy sunflower.

### No evidence for evolution of herbicide resistance

We found no support for evolution of herbicide resistance, as only one individual (< 1 %) of BRW_17_ survived the 1.2 X application rate of imazapyr and no individuals of BRW_07_ survived. At this rate, both homozygous and heterozygous plants carrying the resistant allele should survive (Presotto *et al*. 2012), indicating that the resistant allele is absent, or at a very low frequency in BRW_17_. AHAS resistance is partially dominant, so cultivars with the Clearfield® technology are homozygous for the resistant allele, implying that this allele starts at a high frequency in crop-wild hybrids when the cultivated parent is AHAS resistant. In addition, no fitness costs have been reported in sunflower (White *et al*. 2002). All of this (high initital frequency, dominance, and no fitness costs) makes the rapid loss of the resistant allele due to drift or negative selection very unlikely, and implies that wild and cultivated parents of BRW were susceptible to AHAS herbicides.

Mechanisms of escape to herbicide applications may evolve independently of the evolution of resistance (Darmency *et al*. 2017). For example, delayed seed germination, due to higher seed dormancy, have evolved in many weed species in response to herbicides (Fried *et al*. 2012; Darmency *et al*. 2017). Weed cohorts that emerge later in the season have higher probability of escaping pre-sowing herbicide applications, which could explain how herbicide-susceptible populations may survive and reproduce in maize and sunflower crop fields. On the other hand, in glyphosate-resistant soybean crops, multiple applications during the crop cycle is a common management practice (Singh *et al*. 2020), so emergence should be extremely late to escape herbicide applications, explaining why no weedy plants were observed when soybean was planted (Casquero *et al*. 2013).

### Competitive ability

For competitve ability, we found no differences between BRW_07_ and BRW_17_ under competition with maize, but significant differences were found without competition. In addition, both BRW_07_ and BRW_17_ grew faster than BAR across treatments, suggesting that traits related to rapid growth were already present in the population collected in 2007. Cultivated and wild sunflower largely differ in early growth (Mercer *et al*. 2007; Kost *et al*. 2015), and phenotypic selection analyses suggest that an early rapid growth is advantageous in agricultural environments, therefore, it is expected to be selected rapidly (Kost *et al*. 2015; Presotto *et al*. 2019). Previous studies showed that early rapid growth in BRW_07_ has probably evolved at the expense of lower stress tolerance (Presotto *et al*. 2017), a similar pattern to that observed in North American weedy populations (Mayrose *et al*. 2011). Although we found no differences under competition with maize, the treatments we chose may represent an extreme competition, more typically found in non-agricultural than in agricultural environments (Mercer *et al*. 2007; Presotto *et al*. 2017).

## Conclusions

In this study, we showed that hybridization between cultivated and wild sunflower may lead to the evolution of agricultural weeds, and that these weeds can evolve quickly, sometimes far away from hybrid zones. Moreover, our results provided support for rapid evolution towards higher seed dormancy highlighting the importance of targeting the early stages of establishment for mitigating the impacts of weedy sunflower. To limit the introduction of seeds of sunflower and other weed species to agricultural plots, we recommend the implementation of straightforward management strategies such as cleaning of agricultural machinery. Finally, integrated weed management strategies aimed at preventing the evolution of resistance, such as cover crops, crop rotation, and the use of herbicides with different modes of action, should be a priority in order to avoid the emergence and spread of herbicide resistant populations in Argentina.

## Supplementary material

Supplementary data are available at *Journal of Heredity* online.

## Acknowledgments

We thank CONICET for fellowships for FH and RBV, and Fundación Bunge & Born for a research Fulbright Grant to FH. We also thank Camila Giacomozzi and Ana Julia Laurlund for their valuable assistance with experiments and two anonymous reviewers for their valuable suggestions.

## SUPPLEMENTARY MATERIAL

**Table S1.**
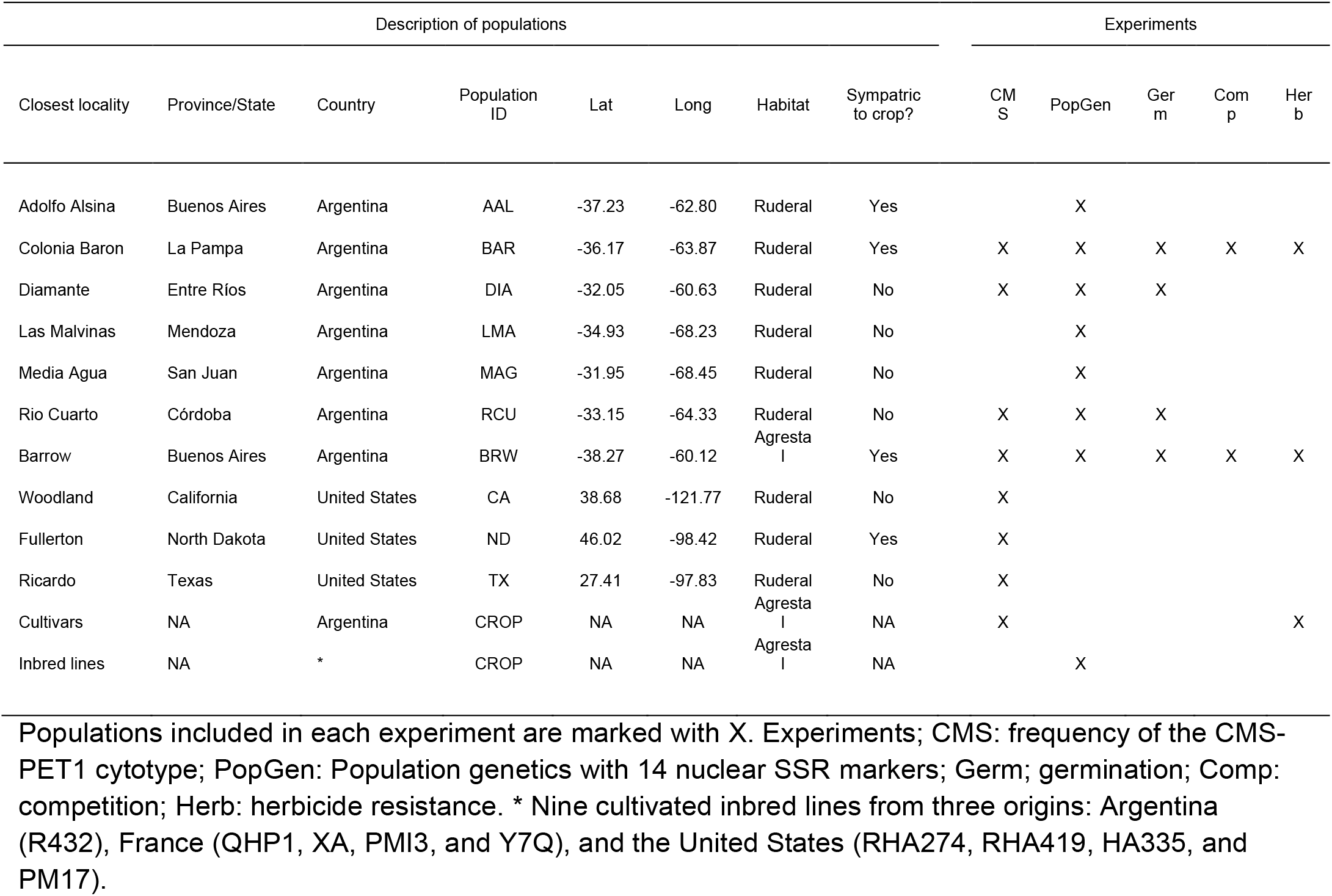
Geographic and habitat information of populations used in this study.

**Fig. S1.**
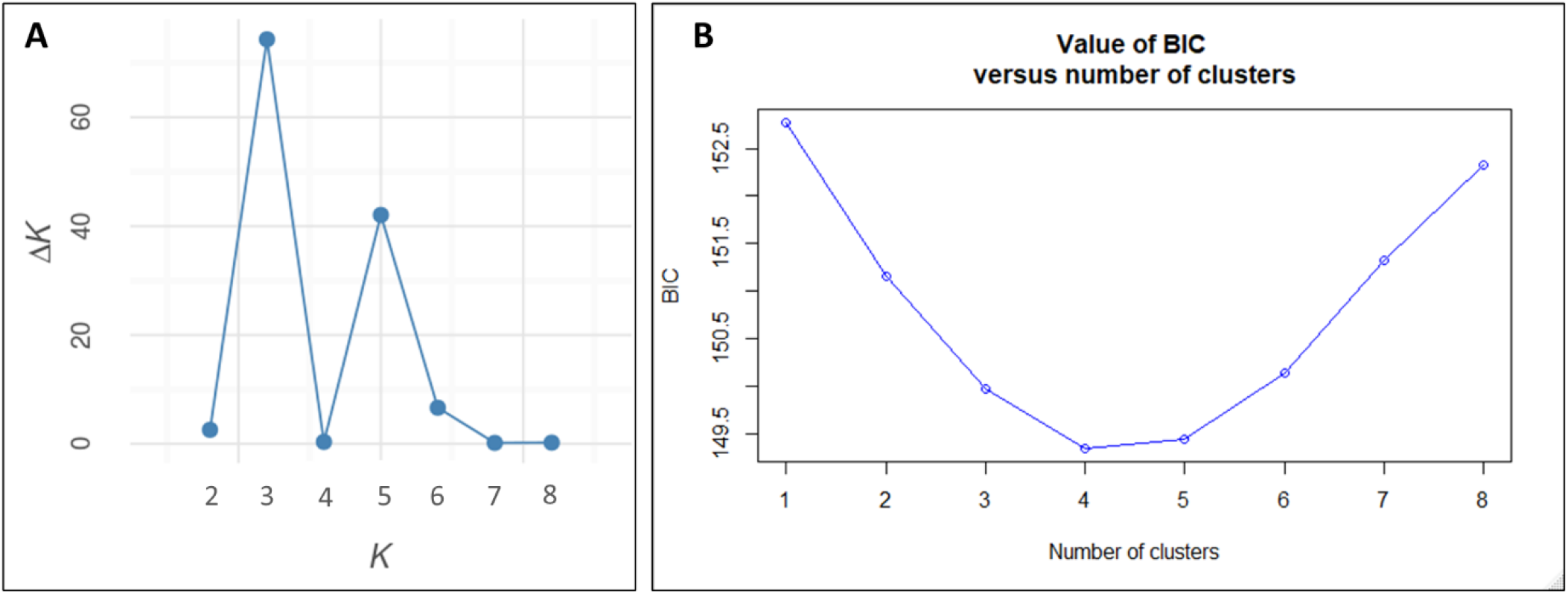
Best supported number of clusters of STRUCTURE (A) and DAPC (B) analyses.

## References

Al-Khatib K, Baumgartner JR, Peterson DE, Currie RS. 1998. Imazethapyr resistance in common sunflower (Helianthus annuus). Weed Science 46: 403–407.

Baker HG. 1974. The Evolution of Weeds. Annual Review of Ecology and Systematics 5: 1–24.

Brunick RL. 2007. Seed dormancy in domesticated and wild sunflowers (Helianthus annuus L.): types, longevity and QTL discovery. PhD Dissertation. Oregon State University, Corvallis, Oregon.

Casquero M, Presotto A, Cantamutto M. 2013. Exoferality in sunflower (Helianthus annuus L.): A case study of intraspecific/interbiotype interference promoted by human activity. Field Crops Research 142: 95–101.

Clement CR. 2014. Landscape Domestication and Archaeology. Encyclopedia of Global Archaeology: 4388–4394.

Le Corre V, Siol M, Vigouroux Y, Tenaillon MI, Délye C. 2020. Adaptive introgression from maize has facilitated the establishment of teosinte as a noxious weed in Europe. Proceedings of the National Academy of Sciences 117: 25618–25627.

Darmency H, Colbach N, Le Corre V. 2017. Relationship between weed dormancy and herbicide rotations: implications in resistance evolution. Pest Management Science 73: 1994– 1999.

Ellstrand NC, Heredia SM, Leak-Garcia JA, et al. 2010. Crops gone wild: Evolution of weeds and invasives from domesticated ancestors. Evolutionary Applications 3: 494–504.

Evanno G, Regnaut S, Goudet J. 2005. Detecting the number of clusters of individuals using the software STRUCTURE: A simulation study. Molecular Ecology 14: 2611–2620.

Filippi CV, Aguirre N, Rivas JG, et al. 2015. Population structure and genetic diversity characterization of a sunflower association mapping population using SSR and SNP markers. BMC Plant Biology 15: 52.

Franks SJ, Hamann E, Weis AE. 2018. Using the resurrection approach to understand contemporary evolution in changing environments. Evolutionary Applications 11: 17–28.

Franks SJ, Kane NC, O’Hara NB, Tittes S, Rest JS. 2016. Rapid genome-wide evolution in Brassica rapa populations following drought revealed by sequencing of ancestral and descendant gene pools. Molecular ecology 25: 3622–3631.

Fried G, Kazakou E, Gaba S. 2012. Trajectories of weed communities explained by traits associated with species’ response to management practices. Agriculture, Ecosystems and Environment 158: 147–155.

Gandhi SD, Heesacker AF, Freeman CA, Argyris J, Bradford K, Knapp SJ. 2005. The self-incompatibility locus (S) and quantitative trait loci for self-pollination and seed dormancy in sunflower. Theoretical and Applied Genetics 111: 619–629.

Garayalde AF, Presotto A, Carrera A, Poverene M, Cantamutto M. 2015. Characterization of a new male sterility source identified in an invasive biotype of Helianthus annuus (L.). Euphytica 206: 579–595.

Goudet J. 2005. HIERFSTAT, a package for R to compute and test hierarchical F-statistics. Molecular Ecology Notes 5: 184–186.

Gressel J. 2005. Introduction – The challenges of ferality. In Gressel J, Crop ferality and volunteerism (pp. 1–7). Boca Raton, FL: CRC Press.

Hernández F, Lindström LI, Parodi E, Poverene M, Presotto A. 2017. The role of domestication and maternal effects on seed traits of crop–wild sunflower hybrids (Helianthus annuus). Annals of Applied Biology 171: 237–251.

Hernández F, Poverene M, Garayalde A, Presotto A. 2019. Re-establishment of latitudinal clines and local adaptation within the invaded area suggest rapid evolution of seed traits in Argentinean sunflower (Helianthus annuus L.). Biological Invasions 21: 2599–2612.

Hernández F, Presotto A, Poverene M, Mandel JR. 2019. Genetic Diversity and Population Structure of Wild Sunflower (Helianthus annuus L.) in Argentina: Reconstructing Its Invasion History. Journal of Heredity 110: 746–759.

Hernández F, Vercellino RB, Fanna I, Presotto A. 2021. Maternal control of early life history traits affects overwinter survival and seedling phenotypes in sunflower (Helianthus annuus L.). Plant Biology 23: 307–316.

Huang Z, Young ND, Reagon M, et al. 2017. All roads lead to weediness: Patterns of genomic divergence reveal extensive recurrent weedy rice origins from South Asian Oryza. Molecular Ecology 26: 3151–3167.

Jombart T, Devillard S, Balloux F. 2010. Discriminant analysis of principal components: a new method for the analysis of genetically structured populations. BMC Genetics 11: 94.

Kane NC, Rieseberg LH. 2008. Genetics and evolution of weedy Helianthus annuus populations: Adaptation of an agricultural weed. Molecular Ecology 17: 384–394.

Kost MA, Alexander HM, Jason Emry D, Mercer KL. 2015. Life history traits and phenotypic selection among sunflower crop-wild hybrids and their wild counterpart: Implications for crop allele introgression. Evolutionary Applications 8: 510–524.

Kreiner JM, Stinchcombe JR, Wright SI. 2018. Population Genomics of Herbicide Resistance: Adaptation via Evolutionary Rescue. Annual Review of Plant Biology 69: 611–635.

Kuester A, Wilson A, Chang SM, Baucom RS. 2016. A resurrection experiment finds evidence of both reduced genetic diversity and potential adaptive evolution in the agricultural weed Ipomoea purpurea. Molecular ecology 25: 4508–4520.

Leclercq P. 1969. Une sterilite male cytoplasmique chez le tournesol. Ann. Amel. Plantes 19: 99–106.

Mayrose M, Kane NC, Mayrose I, Dlugosch KM, Rieseberg LH. 2011. Increased growth in sunflower correlates with reduced defences and altered gene expression in response to biotic and abiotic stress. Molecular Ecology 20: 4683–4694.

Mercer KL, Andow DA, Wyse DL, Shaw RG. 2007. Stress and domestication traits increase the relative fitness of crop-wild hybrids in sunflower. Ecology Letters 10: 383–393.

Mondon A, Owens GL, Poverene M, Cantamutto M, Rieseberg LH. 2018. Gene flow in Argentinian sunflowers as revealed by genotyping-by-sequencing data. Evolutionary Applications 11: 193–204.

Muller MH, Latreille M, Tollon C. 2011. The origin and evolution of a recent agricultural weed: Population genetic diversity of weedy populations of sunflower (Helianthus annuus L.) in Spain and France. Evolutionary Applications 4: 499–514.

Oerke EC. 2006. Crop losses to pests. Journal of Agricultural Science 144: 31–43.

Pandolfo CE, Presotto A, Carbonell FT, Ureta S, Poverene M, Cantamutto M. 2018. Transgene escape and persistence in an agroecosystem: the case of glyphosate-resistant Brassica rapa L. in central Argentina. Environmental Science and Pollution Research 25: 6251– 6264.

Peakall R, Smouse PE. 2012. GenALEx 6.5: Genetic analysis in Excel. Population genetic software for teaching and research-an update. Bioinformatics 28: 2537–2539.

Poverene M, Cantamutto M, Seiler GJ. 2008. Ecological characterization of wild Helianthus annuus and Helianthus petiolaris germplasm in Argentina. Plant Genetic Resources: Characterisation and Utilisation 7: 42–49.

Presotto A, Hernández F, Casquero M, et al. 2020. Seed bank dynamics of an invasive alien species, Helianthus annuus L. Journal of Plant Ecology.

Presotto A, Hernández F, Díaz M, et al. 2017. Crop-wild sunflower hybridization can mediate weediness throughout growth-stress tolerance trade-offs. Agriculture, Ecosystems and Environment 249: 12–21.

Presotto A, Hernández F, Mercer KL. 2019. Phenotypic selection under two contrasting environments in wild sunflower and its crop–wild hybrid. Evolutionary Applications: 1–15.

Presotto A, Ureta MS, Cantamutto M, Poverene M. 2012. Effects of gene flow from IMI resistant sunflower crop to wild Helianthus annuus populations. Agriculture, Ecosystems and Environment 146: 153–161.

Pritchard JK, Stephens M, Donnelly P. 2000. Inference of population structure using multilocus genotype data. Genetics 155: 945–959.

Rieseberg LH, Van Fossen C, Arias D, Carter RL. 1994. Cytoplasmic Male Sterility in Sunflower: Origin, Inheritance, and Frequency in Natural Populations. Journal of Heredity 85: 233–238.

Singh V, Etheredge L, McGinty J, Morgan G, Bagavathiannan M. 2020. First case of glyphosate resistance in weedy sunflower (Helianthus annuus). Pest Management Science 76: 3685–3692.

Suarez-Gonzalez A, Lexer C, Cronk QCB. 2018. Adaptive introgression: a plant perspective. Biology Letters 14: 20170688.

Tan S, Evans RR, Dahmer ML, Singh BK, Shaner DL. 2005. Imidazolinone-tolerant crops: history, current status and future. Pest Management Science 61: 246–257.

Tranel PJ, Wright TR. 2002. Resistance of weeds to ALS-inhibiting herbicides: what have we learned? Weed Science 50: 700–712.

Tranel PJ, Wright TR, Heap IM. 2021. Mutations in herbicide-resistant weeds to ALS inhibitors. Online http://www.weedscience.com. 3/23/2021

Ureta MS, Cantamutto M, Carrera A, Delucchi C, Poverene M. 2008. Natural hybrids between cultivated and wild sunflowers (Helianthus spp.) in Argentina. Genetic Resources and Crop Evolution 55: 1267–1277.

Vercellino RB, Pandolfo CE, Breccia G, Cantamutto M, Presotto A. 2018. AHAS Trp574Leu substitution in Raphanus sativus L.: screening, enzyme activity and fitness cost. Pest Management Science 74: 1600–1607.

Vercellino RB, Pandolfo CE, Cerrota A, Cantamutto M, Presotto A. 2019. The roles of light and pericarp on seed dormancy and germination in feral Raphanus sativus (Brassicaceae). Weed Research 59: 396–406.

Wallace BC, Lajeunesse MJ, Dietz G, et al. 2017. OpenMEE : Intuitive, open-source software for meta-analysis in ecology and evolutionary biology. Methods in Ecology and Evolution 8: 941–947.

Weis AE. 2018. Detecting the “invisible fraction” bias in resurrection experiments. Evolutionary Applications 11: 88–95.

De Wet JMJ, Harlan JR. 1975. Weeds and Domesticates: Evolution in the man-made habitat. Economic Botany 29: 99–108.

White AD, Owen MDK, Hartzler RG, Cardina J. 2002. Common sunflower resistance to acetolactate synthase–inhibiting herbicides. Weed Science 50: 432–437.

